## Supplementary material for "Dynamic modes of Notch transcription hubs conferring memory and stochastic activation revealed by live imaging the co-activator Mastermind"

**De Haro Arbona *et al*: Supplementary Figure Legends and Tables.**

#### Supplementary Figures

Figure S1. Related to Figure 1. Image analysis and effect of Hairless depletion on CSL and Mam.

Figure S2. Related to Figure 2. Notch activation complexes show hub-like properties.

Figure S3. Related to Figure 3. Mediator complex, but not CBP/p300, is involved in Mam enrichment.

Figure S4. Related to Figure 4. Mam perturbations do not affect CSL or IDR enrichment.

Figure S5. Related to Figure 5. Notch deactivation leads to loss of *E(spl)-m3* transcription.

Figure S6. Related to Figure 6. Population fitting reveals infrequent enrichment of Pol II and Med1 but is augmented by ecdysone treatment.

#### Supplementary Tables

Table S1: Genomic co-ordinates and oligonucleotides used for CRISPR, constructs and qPCR.

Table S2: Summary of Drosophila strains.

Table S3: Genetic combinations for each figure

Table S4: p-values from statistical tests.

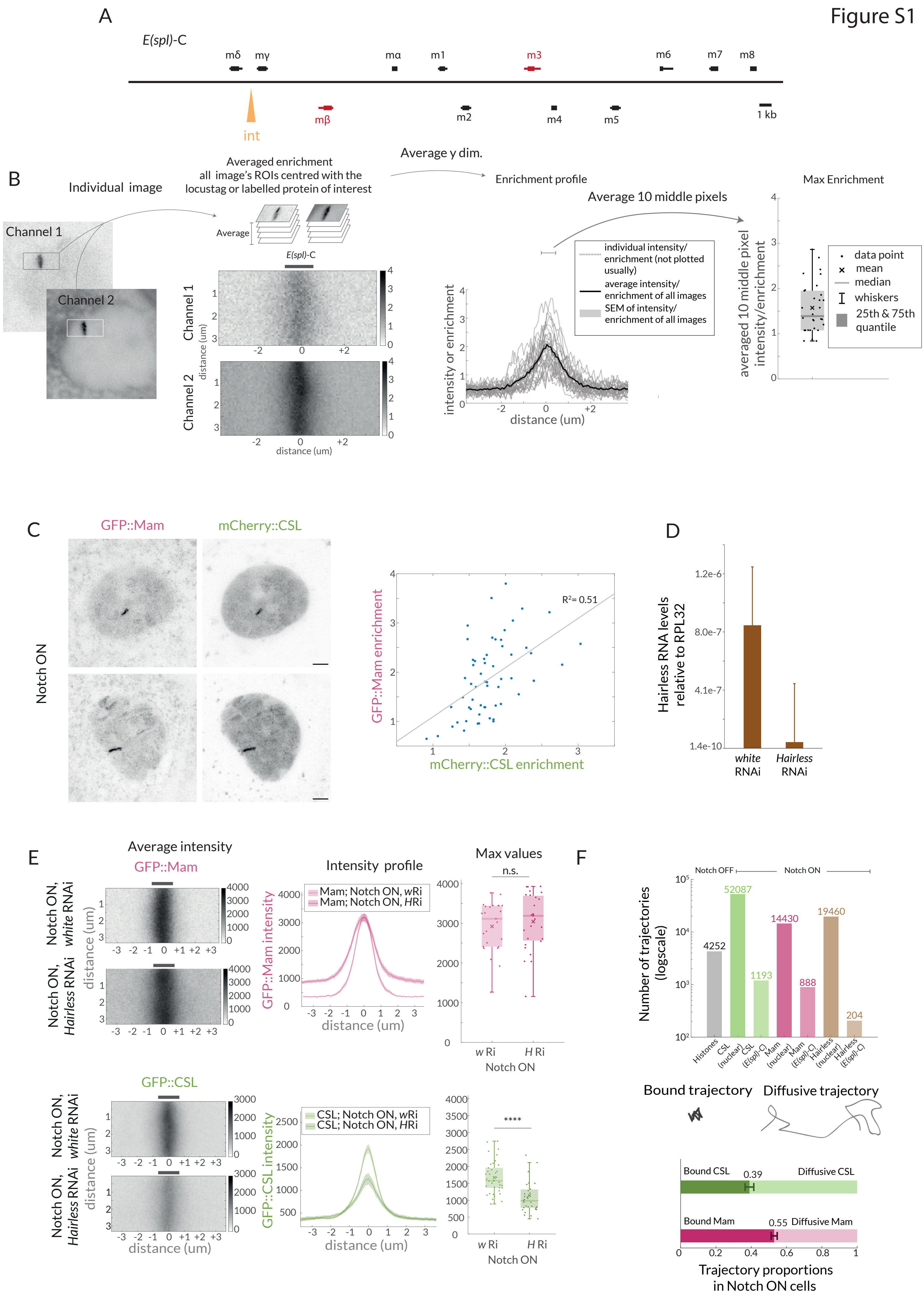
Figure S1
A
E(spl)-C
mδ
mγ
mα
m1
m3
m6
m7
m8
int
mβ
m2
m4
m5
1 kb
B
Individual image
Channel 1
Channel 2
Averaged enrichment
all image's ROIs centred with the
locustag or labelled protein of interest
Average
E(spl)-C
distance (um)
Average y dim.
Enrichment profile
Average 10 middle pixels
individual intensity/
enrichment (not plotted
usually)
average intensity/
enrichment of all images
SEM of intensity/
enrichment of all images
intensity or enrichment
distance (um)
Max Enrichment
averaged 10 middle pixel
intensity/enrichment
data point
mean
median
whiskers
25th & 75th
quantile
C
GFP::Mam
mCherry::CSL
Notch ON
GFP::Mam enrichment
mCherry::CSL enrichment
R²= 0.51
D
Hairless RNA levels
relative to RPL32
1.2e-6
8.0e-7
4.1e-7
1.4e-10
white
RNAi
Hairless
RNAi
E
Average intensity
GFP::Mam
Notch ON, white RNAi
Notch ON, Hairless RNAi
distance (um)
Intensity profile
Mam; Notch ON, wRi
Mam; Notch ON, HRi
GFP::Mam intensity
distance (um)
Max values
n.s.
w Ri
H Ri
Notch ON
GFP::CSL
CSL; Notch ON, wRi
CSL; Notch ON, HRi
GFP::CSL intensity
****
F
Notch OFF
Notch ON
Number of trajectories
(logscale)
4252
52087
1193
14430
888
19460
204
Histones
CSL (nuclear)
CSL (E(spl)-C)
Mam (nuclear)
Mam (E(spl)-C)
Hairless (nuclear)
Hairless (E(spl)-C)
Bound trajectory
Diffusive trajectory
Bound CSL
0.39
Diffusive CSL
Bound Mam
0.55
Diffusive Mam
Trajectory proportions
in Notch ON cells

**Figure S1. Related to Figure 1. Image analysis and effect of Hairless depletion on CSL and Mam.**

(A) Scheme of the *E(spl)*-C in the *Drosophila* genome. Orange arrow indicates the insertion site of the IntB sequence that recruits ParB, *E(spl)-m*$\beta$ and *E(spl)-m*3 are shaded in magenta.

(B) Schematic overview of image analysis. Images were rotated and a ROI was selected centred at the *E(spl)-C*. ROIs from images were averaged to produce an average enrichment image. 2D images extracted with the ROI were averaged in the y dimension to generate an intensity profile enrichment plot, where the mean intensity and SEM were represented. For max values, the 10 middle values of the profile enrichment were averaged and plotted. Box encompasses range between 0.25 and 0.75 quantile, whiskers extend to furthest points not considered outliers, bar marks median, cross marks the mean and each dot is the value for one nucleus.

(C) Two colour images of Mam and CSL (labelled with GFP and mCherry, respectively) in Notch ON cells. Graph depicts correlation of the enrichment levels.

(D) mRNA levels of *Hairless* in control (*white RNAi*) and *Hairless RNAi* conditions measured by RT-qPCR and normalised to RPL32.

(E) Average enrichment, intensity profile and max values for Mam (n = 25, 27) and CSL (n = 37, 31) in *Hairless* RNAi (H Ri) in comparison to control, *white* RNAi (*w* Ri). For p-see Table S4.

(F) Graph shows number of trajectories for each molecule, all but histones are Notch ON and separated in nuclear and at the *E(spl)-C*. Trajectories were segregated into bound and diffusive, as depicted by the cartoon. Percentages of each population for CSL and Mam were calculated per nuclei, error represents SEM (Percentage CSL vs. Mam p = 0.003, n= 7, 13 nuclei). Diffusion coefficients were calculated per nucleus as well. Diffusion coefficient bound CSL vs. bound Mam (0.27 ± 0.04 um2/s, 0.25 ± 0.01 um2/s p = 0.63), Diffusion coefficient diffusive CSL vs. bound Mam (0.01 ± 0.001 um2/s, 0.01 ± 0.001 um2/s, p = 0.34).

Figure S2

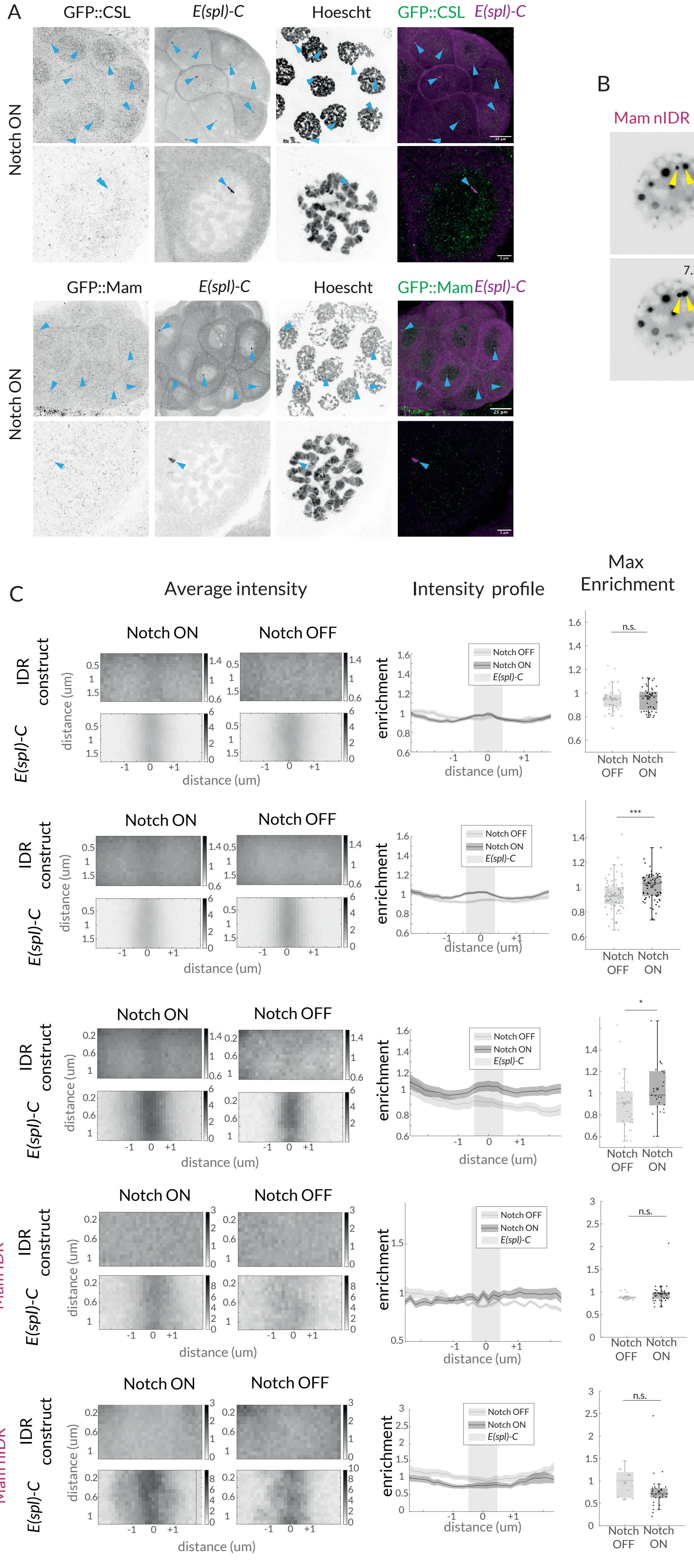
A
GFP::CSL
E(spl)-C
Hoescht
GFP::CSL E(spl)-C
Notch ON
GFP::Mam
E(spl)-C
Hoescht
GFP::Mam E(spl)-C
Notch ON
B
Mam nIDR
0 sec
2.58 sec
5.16 sec
7.74 sec
10.32 sec
12.9 sec
5 µm
C
Average intensity
Intensity profile
Max Enrichment
Notch ON
Notch OFF
CSL ct IDR
NICD IDR
Med1 IDR
Mam IDR
Mam nIDR
IDR construct
E(spl)-C
distance (um)
enrichment
Notch OFF
Notch ON
n.s.
***
*
Notch OFF
Notch ON

**Figure S2. Related to Figure 2. Notch activation complexes exhibit hub-like properties**

(A) Fixed Notch ON whole glands and nuclei of GFP::CSL and GFP::Mam (first channel), co-stained to detect *E(spl)*-C locus and DNA (Hoechst). Top row, whole gland and lower row individual example nuclei, with 25um and 5um scale bars respectively

(B) An example of a nucleus expressing GFP::Mam[nIDR], imaged over time. Yellow arrows mark two droplets that fuse (10 sec frame). Scale bar indicates 5um.

(C) Average, profile and max enrichment of IDR constructs: CSL[ctIDR], NICD[IDR], Med1[IDR], Mam, Mam[nIDR] in Notch OFF and Notch ON cells (upper and lower rows). For p-values see Table S4.

### Figure S3

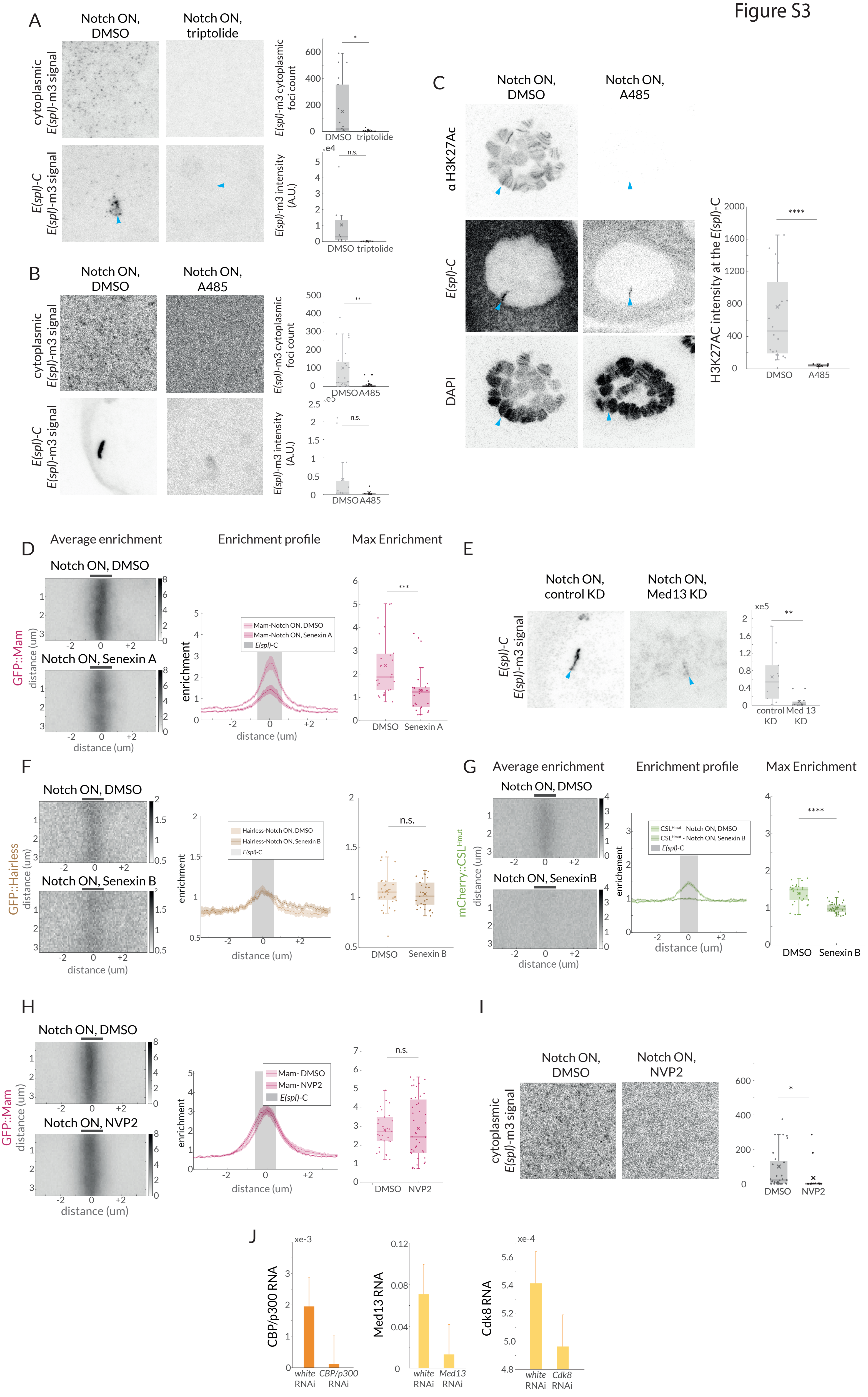
A
Notch ON, DMSO
Notch ON, triptolide
cytoplasmic E(spl)-m3 signal
E(spl)-C E(spl)-m3 signal
E(spl)-m3 cytoplasmic foci count
DMSO triptolide
E(spl)-m3 intensity (A.U.)
n.s.
B
Notch ON, DMSO
Notch ON, A485
E(spl)-m3 cytoplasmic foci count
E(spl)-m3 intensity (A.U.)
DMSO A485
C
Notch ON, DMSO
Notch ON, A485
α H3K27Ac
E(spl)-C
DAPI
H3K27AC intensity at the E(spl)-C
****
D
Average enrichment
Enrichment profile
Max Enrichment
Notch ON, DMSO
Notch ON, Senexin A
GFP::Mam
distance (um)
enrichment
Mam-Notch ON, DMSO
Mam-Notch ON, Senexin A
E(spl)-C
DMSO Senexin A
E
Notch ON, control KD
Notch ON, Med13 KD
E(spl)-C E(spl)-m3 signal
control KD
Med 13 KD
F
Notch ON, DMSO
Notch ON, Senexin B
GFP::Hairless
Hairless-Notch ON, DMSO
Hairless-Notch ON, Senexin B
DMSO Senexin B
G
Average enrichment
Enrichment profile
Max Enrichment
Notch ON, DMSO
Notch ON, SenexinB
mCherry::CSL Hmut
CSLHmut - Notch ON, DMSO
CSLHmut - Notch ON, Senexin B
DMSO Senexin B
H
Notch ON, DMSO
Notch ON, NVP2
GFP::Mam
Mam- DMSO
Mam- NVP2
DMSO NVP2
I
Notch ON, DMSO
Notch ON, NVP2
cytoplasmic E(spl)-m3 signal
DMSO NVP2
J
CBP/p300 RNA
white RNAi
CBP/p300 RNAi
Med13 RNA
Med13 RNAi
Cdk8 RNA
Cdk8 RNAi

**Figure S3. Related to Figure 3. Mediator complex, but not CBP/p300, is involved in Mam enrichment.**

(A) Expression of *E(spl)*-m3, detected by smFISH, is lost upon triptolide treatment in Notch ON cells. Representative images show number of cytoplasmic RNA puncta in the volume sampled (n = 18, 16 for DMSO, triptolide) and intensity at the *E(spl)-C* (indicated by blue arrow for triptolide, n = 9, 8 for DMSO, triptolide). p-values provided in Table S4.

(B) Expression of *E(spl)*-m3, detected by smFISH, is lost upon A485 treatment in Notch ON cells. Representative images show number of cytoplasmic RNA puncta in the volume sampled (n = 31, 22 for DMSO, A485) and intensity at the *E(spl)-C* (n = 13, 6 for DMSO, A485).

(C) Representative images of H3K27Ac staining in Notch ON glands treated with DMSO and A485. H3K27Ac (top row) is lost throughout the nucleus and at *E(spl)-C* (blue arrows, middle row). Quantification of H3K27Ac levels at *E(spl)-C* (n = 19, 6 for DMSO and A485). p-values provided in Table S4.

(D) Average, profile and max enrichment of Mam in Notch ON cells, treated with DMSO or Senexin A. (n = 27, 26 for DMSO and Senexin). p-values provided in Table S4.

(E) Expression of *E(spl)*-m3, detected by smFISH, is lost in the Med13 KD in Notch ON cells compared to control KD (white RNAi). Representative images from the nucleus (centred on *E(spl)-C*, *blue* arrow) and quantification of intensity with boxplots, as described in Figure 1B (n = 10, 11 for control KD and Med13 KD).

(F) Average, profile, and max enrichment of Hairless in Notch ON cells, treated with DMSO or Senexin B. (n = 31, 28  for DMSO and Senexin B). All p-values provided in Table S4.

(G) Average, profile, and max enrichment of $CSL^{Hmut}$ in Notch ON nuclei, treated with DMSO or Senexin B. (n = 27, 31 for DMSO and Senexin B). All p-values provided in Table S4.

(H) Average, profile, and max enrichment of Mam in Notch ON nuclei, treated with DMSO or NVP2. (n = 30, 43for DMSO and NVP2).

(I) Expression of *E(spl*-m3, detected by smFISH, is lost upon NVP2 treatment in Notch ON cells. Representative images illustrate density of RNA puncta. (n = 31, 14 for DMSO and NVP2). All p-values provided in Table S4.

(J) mRNA levels of *nej* (CBP/p300), *Med13* and *Cdk8* in tissues exposed to the respective RNAi or to control RNAi, measured by RT-qPCR and normalised to RPL32.

Figure S4

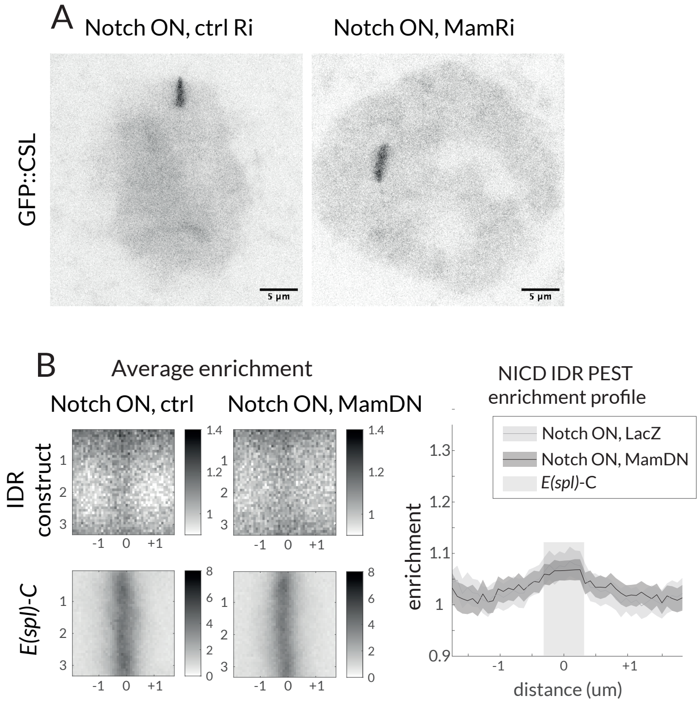
A
Notch ON, ctrl Ri
Notch ON, MamRi
GFP::CSL
5 µm
5 µm
B
Average enrichment
Notch ON, ctrl
Notch ON, MamDN
IDR construct
E(spl)-C
NICD IDR PEST enrichment profile
Notch ON, LacZ
Notch ON, MamDN
E(spl)-C
enrichment
distance (um)

**Figure S4. Related to Figure 4. Mam perturbations do not affect CSL or IDR enrichment**

(A) Representative GFP::CSL images at the *E(spl)-C* in Notch ON nuclei exposed to ctrl (*y-RNAi*) or *Mam-RNAi*. Scale bars represent 5um.

(B) GFP::NICD[PEST-IDR] average enrichment and profile in Notch ON nuclei exposed to control (lacZ) or Mam[DN] expression. Images were centred on *E(spl)-C*, lower left(n = 25, 28 for ctrl and Mam[DN)]. p-value provided in Table S4.

### Figure S5

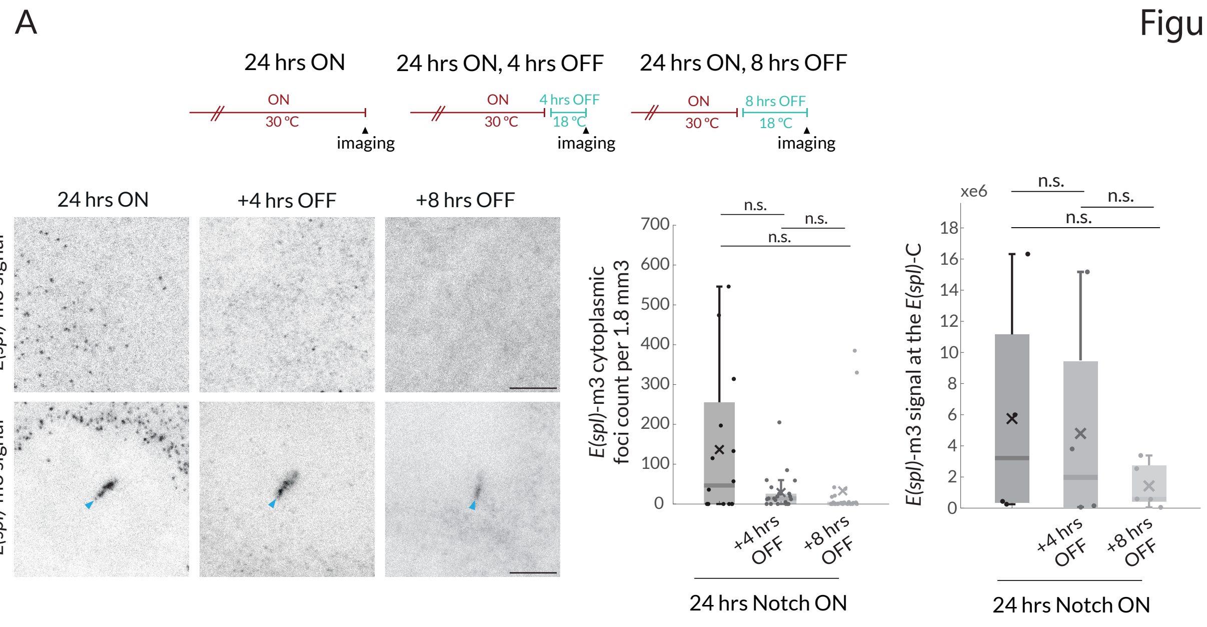
A
24 hrs ON
24 hrs ON, 4 hrs OFF
24 hrs ON, 8 hrs OFF
ON
30 °C
imaging
4 hrs OFF
18 °C
8 hrs OFF
24 hrs ON
+4 hrs OFF
+8 hrs OFF
cytoplasmic E(spl)-m3 signal
E(spl)-C E(spl)-m3 signal
n.s.
E(spl)-m3 cytoplasmic foci count per 1.8 mm3
0
100
200
300
400
500
600
700
+4 hrs OFF
+8 hrs OFF
24 hrs Notch ON
xe6
E(spl)-m3 signal at the E(spl)-C
0
2
4
6
8
10
12
14
16
18
24 hrs Notch ON

**Figure S5. Related to Figure 4. Notch deactivation leads to loss of *E(spl)*-m3 transcription.**

(A) Cartoon depicts temperature paradigm, similar to Figure 5A. Expression of *E(spl)*-m3 in the conditions indicated, detected by smFISH. Upper panels show representative cytoplasmic sections (1.8 $mm^3$) and lower signal at the *E(spl)*-C. (n = 22, 30,30) for cytoplasmic 24 hrs ON, 24 hrs ON + 4 hrs OFF and 24 hrs ON + 8 hrs OFF; n = 4, 4, 4 for nuclear 24 hrs ON, 24 hrs ON + 4 hrs OFF and 24 hrs ON + 8 hrs OFF). p-values provided in Table S4.

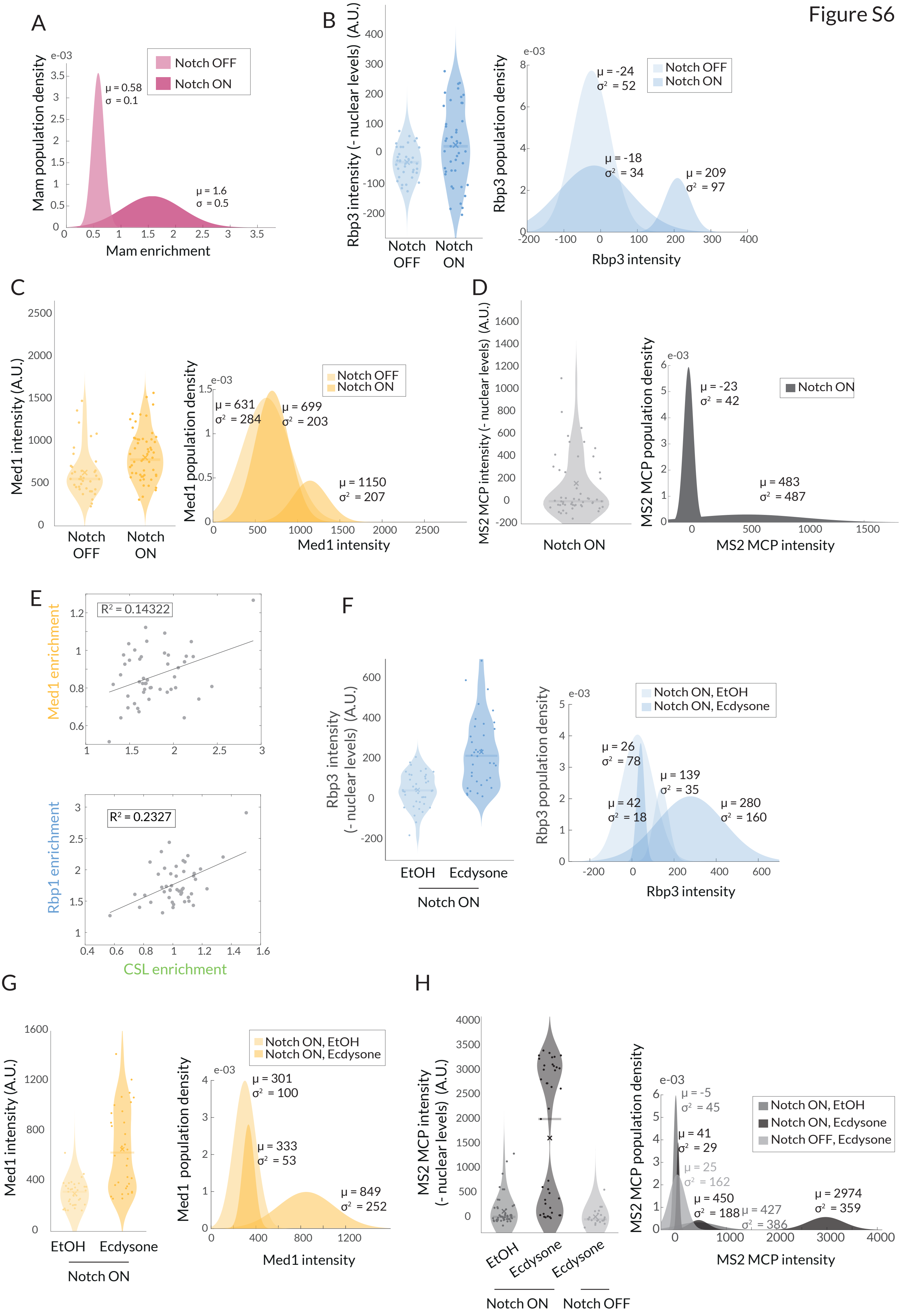

Figure S6
A
Mam population density
Notch OFF
Notch ON
μ = 0.58
σ = 0.1
μ = 1.6
σ = 0.5
Mam enrichment
B
Rbp3 intensity (- nuclear levels) (A.U.)
Notch OFF
Notch ON
Rbp3 population density
μ = -24
σ² = 52
μ = -18
σ² = 34
μ = 209
σ² = 97
Rbp3 intensity
C
Med1 intensity (A.U.)
Notch OFF
Notch ON
Med1 population density
μ = 631
σ² = 284
μ = 699
σ² = 203
μ = 1150
σ² = 207
Med1 intensity
D
MS2 MCP intensity (- nuclear levels) (A.U.)
Notch ON
MS2 MCP population density
μ = -23
σ² = 42
μ = 483
σ² = 487
MS2 MCP intensity
E
Med1 enrichment
R² = 0.14322
Rbp1 enrichment
R² = 0.2327
CSL enrichment
F
Rbp3 intensity (- nuclear levels) (A.U.)
EtOH
Ecdysone
Notch ON
Rbp3 population density
Notch ON, EtOH
Notch ON, Ecdysone
μ = 26
σ² = 78
μ = 42
σ² = 18
μ = 139
σ² = 35
μ = 280
σ² = 160
Rbp3 intensity
G
Med1 intensity (A.U.)
EtOH
Ecdysone
Notch ON
Med1 population density
Notch ON, EtOH
Notch ON, Ecdysone
μ = 301
σ² = 100
μ = 333
σ² = 53
μ = 849
σ² = 252
Med1 intensity
H
MS2 MCP intensity (- nuclear levels) (A.U.)
EtOH
Ecdysone
Ecdysone
Notch ON
Notch OFF
MS2 MCP population density
Notch ON, EtOH
Notch ON, Ecdysone
Notch OFF, Ecdysone
μ = -5
σ² = 45
μ = 41
σ² = 29
μ = 25
σ² = 162
μ = 450
σ² = 188
μ = 427
σ² = 386
μ = 2974
σ² = 359
MS2 MCP intensity

**Figure S6. Related to Figure 6. Population fitting reveals infrequent enrichment of Pol II and Med1 but is augmented by ecdysone treatment.**

(A-D, F-H) Gaussian fitting of Mam enrichment data from Figure 1 and of Rbp3, Med1 and MS2 in the indicated conditions. Violin plots show the distribution of the data with dots depicting intensity or enrichment per nucleus, x average and bar median. μ indicates population average and σ the standard deviation.

(E) Correlation of GFP::Med1 and Halo::CSL, mCherry::Rbp1 and Halo::CSL.

**Supplementary Tables**

**Table S1: Genomic co-ordinates and oligonucleotides used for CRISPR, constructs and qPCR.**

| | |
|---|---|
| Mam gRNA | ATGAATATTTGGGTAAGATT |
| Mam homology arm upstream | 2R:14009456..14010278 |
| Mam homology arm downstream | 2R:14009456..14010278 |
| Med1 gRNA | TAGGAACGCATAGATATGAG |
| Med1 homology arm upstream | 3L:21633297..21634295 |
| Med1 homology arm downstream | 3L:21634297..21635295 |
| *E(spl)mβ HLH* gRNA | GGGACGCCACATGGGGCCAG |
| *E(spl)mβ HLH* homology arm upstream | 3R:26005336..26006118 |
| *E(spl)mβ-HLH* homology arm downstream | 3R:26004521..26005328 |
| CSL nt IDR | 2L:15039938..15040106 |
| CSL ct IDR | 2L:15042401..15042618 |
| NICD IDR | X: 3170337..3170534 |
| NICD PEST IDR | X: 3169065..3170534 |
| Med1 IDR | 3L:21629087..21632060 |
| Mam IDR | 2R:14055747..14057930 |
| Mam nIDR | 2R:14026225..14064332 |
| cDNA: RPL32 fwd | ATGCTAAGCTGTCGCACAAATG |
| cDNA: RPL32 rvs | GTTCGATCCGTAACCGATGT |
| cDNA: Hairless fwd | CATCGCTGAGCTTTTCGGAC |
| cDNA: Hairless rvs | CTCGGCCAAGCGTACTGTTC |
| cDNA: CBP/p300 fwd | AACGCCATACCAGGCATGAA |
| cDNA: CBP/p300 rvs | GAATTGATCATCCCACCGCC |
| cDNA: Med13 fwd | TGGAAGTGGTGTCGGGTGGT |
| cDNA: Med13 rvs | GCCAGAACAGCTGACTCGGC |
| cDNA: Cdk8 fwd | AGATGAAAACGCAGATAGAGCG |
| cDNA: Cdk8 rvs | TTCTTTGCCATCGCTTGTCT |

**Table S2: Summary of Drosophila strains.**

| Short name | Notes | Source | RRID, reference. |
|---|---|---|---|
| 1151-Gal4 | | | FBti0007229 |
| vas-phiC31;;M [attP86Fb] | Used for 3rd chromosome insertions | Bloomington Stock Center | RRID:BDSC_24749 |
| UAS-N∆ECD | 2nd and 3rd chromosome insertions (unknown and attP86Fb insertion) | [37,38] | |
| eGFP::CSL | Genomic fragment insertion attP86Fb (RRID:BDSC 24749) | [34] | |
| Halo::CSL | Genomic fragment insertion attP86Fb | [69] | |
| vas-phiC31;;M [attP51D] | Used for 2nd chromosome insertions | | RRID:BDSC 24483 |
| eGFP::Hairless | Genomic fragment insertion attP51D | [34] | |
| Hairless::Halo | Genomic fragment insertion attP51D | [42] | |
| Med13::YFP | | Bloomington Stock Center | RRID:BDSC_57899 |
| UAS-ParA1-mCherry | Insertion attP86Fb | [34] | |
| UAS-ParB1-GFP | Insertion attP86Fb | [34] | |
| *E(spl)-C[*m∂ intA] | CRISPR of E(spl)-C | [34] | |
| UAS-Hairless-RNAi | | Bloomington Stock Center; Line used in: [34] | RRID:BDSC_27315 |
| UAS-Mam-RNAi | | Bloomington Stock Center; Line used in: [34,108,109] | RRID:BDSC_28046 |
| UAS-Med13-RNAi | | Bloomington Stock Center; Line used in: [110,111] | RRID:BDSC_34630 |
| UAS-nejire-RNAi | | Vienna Resource Center | KK10288 |
| UAS-Cdk8-RNAi | | [61] | |
| UAS-Mam[DN] | | [67] | |
| MS1096-Gal4 | | Bloomington Stock Center | RRID:BDSC_8860 |

| | | | |
|---|---|---|---|
| nanos-phiC31; Msp300[attP40] | Used for 2nd chromosome insertions of UAS-IDR | Bloomington Stock Center | RRID:BDSC_25709 |
| UAS-Cry2-TevC | Insertion AttP51C (RRID:BDSC 24482) | [69] | |
| UAS-OptIC-Notch{ω}[mCherry] | Insertion attP40 | [69] | |
| *E(spl)-mβ*-HLH-MS2-LacZ | CRISPR of E(spl)-*mβ* | This paper, modified from [11] | |
| hsp83-MCP::GFP | | Bloomington Stock Center | RRID:BDSC_7280 |
| nos-cas9 | | Bloomington Stock Center | RRID:BDSC_54591 |

| | | | |
|---|---|---|---|
| αtub-piggyBac | | Bloomington Stock Center | RRID:BDSC_32070 |
| sfGFP::Mam | CRISPR of Mam locus | This paper | |
| Halo::Mam | CRISPR of Mam locus | This paper | |
| H2AV::Halo | | [42] | |
| sfGFP::Med1 | CRISPR of Med1 locus | This paper | |
| eGFP::Rbp3 | CRISPR of Rbp3 locus | [16] | |
| mCherry::Rbp1 | CRISPR of Rbp3 locus | [16] | |
| UAS-sfGFP-NICD[IDR] | Insertion attp40 | This paper | |
| UAS-sfGFP-NICD[IDRPEST] | Insertion attp40 | This paper | |
| UAS-sfGFP-CSL[ntIDR] | Insertion attp40 | This paper | |
| UAS-sfGFP-CSL[ctIDR] | Insertion attp40 | This paper | |
| UAS-sfGFP-CSL[ctIDR] | Insertion attp40 | This paper | |
| UAS-sfGFP-Med1[IDR] | Insertion attp40 | This paper | |
| UAS-sfGFP-Mam[nIDR] | Insertion attp40 | This paper | |
| UAS-sfGFP-Mam[IDR] | Insertion attp40 | This paper | |

**Table S3: Genetic combinations used for each Figure**

| Figure | X chromosome | II chromosome | III chromosome |
|---|---|---|---|
| Figure 1B | *1151-Gal4* | *UAS-NΔECD, GFP::Mam* | *E(spl)-C[m∂ intA}, UAS-ParA::mCherry* |
| Figure 1B | *1151-Gal4* | *UAS-NΔECD* | *GFP::CSL, E(spl)-C[m∂ intA}, UAS-ParA::mCherry* |
| Figure 1C | *1151-Gal4* | *UAS-NΔECD, GFP::Mam* | *UAS-white RNAi* |
| Figure 1C | *1151-Gal4* | *UAS-NΔECD* | *UAS-white RNA, GFP::CSL* |
| Figure 1C | *1151-Gal4* | *UAS-NΔECD, GFP::Mam* | *UAS-Hairless RNAi* |
| Figure 1C | *1151-Gal4* | *UAS-NΔECD, GFP::Hairless* | *E(spl)-C[m∂ intA}, UAS-ParA::mCherry* |
| Figure 1C' | *1151-Gal4* | *UAS-NΔECD* | *UAS-Hairless RNAi, GFP::CSL* |
| Figure 1C' | *1151-Gal4* | *UAS-NΔECD* | *UAS-white-RNAi, GFP::CSL* |

| | | | |
|---|---|---|---|
| Figure 1D | *1151-Gal4* | *UAS-LacZ* | *E(spl)-C[m∂ intA}, UAS-ParB::GFP*, *Halo::CSL* |
| Figure 1D | *1151-Gal4* | *UAS-NΔECD* | *E(spl)-C[m∂ intA}, UAS-ParB::GFP*, *Halo::CSL* |
| Figure 1D | *1151-Gal4* | *UAS-LacZ, Halo::Mam* | *E(spl)-C[m∂ intA}, UAS-ParB::GFP* |
| Figure 1D | *1151-Gal4* | *UAS-NΔECD, Halo::Mam* | *E(spl)-C[m∂ intA}, UAS-ParB::GFP* |
| Figure 1E-E' | *1151-Gal4* | *UAS-NΔECD* | *E(spl)-C[m∂ intA}, UAS-ParB::GFP*, *Halo::CSL* |
| Figure 1E-E' | *1151-Gal4* | *UAS-NΔECD, Halo::Mam* | *E(spl)-C[m∂ intA}, UAS-ParB::GFP* |
| Figure 1E-E' | *1151-Gal4* | *UAS-NΔECD, Halo ::Hairless* | *E(spl)-C[m∂ intA}, UAS-ParA::mCherry* |
| Figure S1C | *ms1096-Gal4, tub-Gal80ts* | *UAS-NΔECD, GFP::Mam* | *mCherry::CSL* |
| Figure S1D | *1151-Gal4* | *UAS-NΔECD* | *UAS-white RNAi* |
| Figure S1D | *1151-Gal4* | *UAS-NΔECD* | *UAS-Hairless RNAi* |
| Figure S1E | *As Figure 1C* | | |
| Figure S1F | *As Figure 1D-E'* | | |
| Figure 2A | *1151-Gal4* | *UAS-NΔECD* | *GFP::CSL, 12xCSLsites* |
| Figure 2A | *1151-Gal4* | *UAS-NΔECD* | *GFP::CSL, 48xCSLsites* |
| Figure 2C | *ms1096-Gal4, tub-Gal80ts* | *UAS-NΔECD, UAS-NICD[PEST IDR]::GFP* | *E(spl)-C[m∂ intA}, UAS-ParA::mCherry* |

| | | | |
|---|---|---|---|
| Figure 2C | *ms1096-Gal4, tub-Gal80ts* | *UAS-NΔECD, UAS-CSL[ntIDR]::GFP* | *E(spl)-C[m∂ intA}, UAS-ParA::mCherry* |
| Figure 2D | *1151-Gal4* | *UAS-NΔECD, GFP::Mam* | *E(spl)-C[m∂ intA}, UAS-ParA::mCherry* |
| Figure 2D | *1151-Gal4* | *UAS-NΔECDΔIDR, GFP::Mam* | *E(spl)-C[m∂ int A}, UAS-ParA::mCherry* |
| Figure 2E | *1151-Gal4* | *UAS-NΔECD, Halo::Mam* | *E(spl)-C[m∂ intA}, UAS-ParB::GFP* |
| Figure 2E | *1151-Gal4* | *UAS-NΔECD* | *E(spl)-C[m∂ intA}, UAS-ParB::GFP*, Halo::CSL |
| Figure S2A | *1151-Gal4* | *UAS-NΔECD* | *E(spl)-C[m∂ intA}, UAS-ParA::mCherry*, GFP::CSL |
| Figure S2A | *1151-Gal4* | *UAS-NΔECD, GFP::Mam* | *E(spl)-C[m∂ intA}, UAS-ParA::mCherry* |
| Figure S2B | *ms1096-Gal4, tub-Gal80ts* | *UAS-NΔECD, UAS-Mam[nIDR]::GFP* | *E(spl)-C[m∂ intA}, UAS-ParA::mCherry* |
| Figure S2C | *ms1096-Gal4, tub-Gal80ts* | *UAS-NΔECD, UAS-CSL[ctIDR]:: GFP* | *E(spl)-C[m∂ intA}, UAS-ParA::mCherry* |
| Figure S2C | *ms1096-Gal4, tub-Gal80ts* | *UAS-NΔECD, UAS-NICD[IDR]:: GFP* | *E(spl)-C[m∂ intA}, UAS-ParA::mCherry* |
| Figure S2C | *ms1096-Gal4, tub-Gal80ts* | *UAS-NΔECD, UAS-Med1[IDR]-GFP* | *E(spl)-C[m∂ intA}, UAS-ParA::mCherry* |
| Figure S2C | *ms1096-Gal4, tub-Gal80ts* | *UAS-NΔECD, UAS-Mam[IDR]-GFP* | *E(spl)-C[m∂ intA}, UAS-ParA::mCherry* |

| | | | |
|---|---|---|---|
| Figure S2C | *ms1096-Gal4, tub-Gal80ts* | *UAS-NΔECD, UAS-Mam[nIDR]-GFP* | *E(spl)-C[m∂ intA}, UAS-ParA::mCherry* |
| Figure 3A-C | *1151-Gal4* | *UAS-NΔECD, GFP::Mam* | *E(spl)-C[m∂ intA}, UAS-ParA::mCherry* |
| Figure 3D | *1151-Gal4* | *GFP::Mam, UAS-yellow-RNAi* | *UAS-NΔECD* |
| Figure 3D | *1151-Gal4* | *GFP::Mam, UAS-nej-RNAi* | *UAS-NΔECD* |
| Figure 3E | *1151-Gal4* | *UAS-NΔECD, GFP::Mam* | *UAS-white-RNAi* |
| Figure 3E | *1151-Gal4* | *UAS-NΔECD, GFP::Mam* | *UAS-med13-RNAi* |
| Figure 3E | *1151-Gal4* | *UAS-NΔECD* | *UAS-white-RNAi, GFP::CSL* |
| Figure 3E | *1151-Gal4* | *UAS-NΔECD* | *UAS-med13-RNAi, GFP::CSL* |
| Figure 3E | *1151-Gal4* | *UAS-NΔECD, GFP::Mam* | *UAS-med13-RNAi* |
| Figure 3F | *1151-Gal4* | *UAS-NΔECD, GFP::Mam* | *E(spl)-C[m∂ intA}, UAS-ParA::mCherry* |
| Figure 3F | *1151-Gal4* | *UAS-NΔECD* | *E(spl)-C[m∂ intA}, UAS-ParA::mCherry, GFP::CSL* |
| Figure 3G | *1151-Gal4* | *UAS-NΔECD, GFP::Mam* | *UAS-white-RNAi* |
| Figure 3G | *1151-Gal4* | *UAS-NΔECD, GFP::Mam* | *UAS-cdk8-RNAi* |
| Figure 3H | *1151-Gal4* | *UAS-NΔECD* | *E(spl)-C[m∂ intA}, UAS-ParA::mCherry, skd(Med13)::YFP* |
| Figure 3H | *1151-Gal4* | *UAS-LacZ* | *E(spl)-C[m∂ intA}, UAS-ParA::mCherry, skd(Med13)::YFP* |
| Figure S3A | *1151-Gal4* | *UAS-NΔECD, GFP::Mam* | *E(spl)-C[m∂ intA}, UAS-ParA::mCherry* |
| Figure S3A-D | *1151-Gal4* | *UAS-NΔECD, GFP::Mam* | *E(spl)-C[m∂ intA}, UAS-ParA::mCherry* |
| Figure S3E | *1151-Gal4* | *UAS-NΔECD, GFP::Mam* | *UAS-white-RNAi* |
| Figure S3E | *1151-Gal4* | *UAS-NΔECD, GFP::Mam* | *UAS-med13-RNAi* |
| Figure S3E | *1151-Gal4* | *UAS-NΔECD, GFP::Mam* | *UAS-white-RNAi* |
| Figure S3F | *1151-Gal4* | *UAS-NΔECD, GFP::Hairless* | *E(spl)-C[m∂ intA}, UAS-ParA::mCherry* |
| Figure S3G | *1151-Gal4* | *UAS-NΔECD, mCherry::CSL[LLL]* | *E(spl)-C[m∂ intA}, UAS-ParA::GFP* |
| Figure S3H, I | *1151-Gal4* | *UAS-NΔECD, GFP::Mam* | *E(spl)-C[m∂ intA}, UAS-ParA::mCherry* |
| Figure S3J | *1151-Gal4* | *UAS-NΔECD* | *UAS-white-RNAi* |
| Figure S3J | *1151-Gal4* | *UAS-NΔECD, UAS CBP/p300 RNAi* | |
| Figure S3J | *1151-Gal4* | *UAS-NΔECD* | *UAS-med13-RNAi* |
| Figure S3J | *1151-Gal4* | *UAS-NΔECD* | *UAS-cdk8-RNAi* |
| Figure 4A | *1151-Gal4* | *GFP::Mam, UAS-yellow-RNAi* | *UAS-NΔECD* |

| | | | |
|---|---|---|---|
| Figure 4A | *1151-Gal4* | *GFP::Mam, UAS-Mam[DN]* | *UAS-NΔECD* |
| Figure 4A' | *1151-Gal4* | *UAS-NΔECD, UAS-LacZ* | *GFP::CSL* |
| Figure 4A' | *1151-Gal4* | *UAS-NΔECD, UAS-Mam[DN]* | *GFP::CSL* |
| Figure 4B | *1151-Gal4* | *UAS LacZ, UAS-LacZ* | |
| Figure 4B | *1151-Gal4* | *UAS-NΔECD, UAS-LacZ* | |
| Figure 4B | *1151-Gal4* | *UAS LacZ, UAS-Mam[DN]* | |
| Figure 4B | *1151-Gal4* | *UAS-NΔECD, UAS-Mam[DN]* | |
| Figure 4C | *1151-Gal4* | *UAS-NΔECD, UAS LacZ* | *E(spl)-C[m∂ intA}, UAS-ParA::mCherry, skd::YFP* |
| Figure 4C | *1151-Gal4* | *UAS-NΔECD, UAS-Mam[DN]* | *E(spl)-C[m∂ intA}, UAS-ParA::mCherry, skd::YFP* |
| Figure 4D-D' | *1151-Gal4* | *UAS-NΔECD, UAS-LacZ* | |
| Figure 4D-D' | *1151-Gal4* | *UAS LacZ, UAS-Mam[DN]* | |
| Figure 4D-D' | *1151-Gal4* | *UAS-NΔECD, UAS-Mam[DN]* | |
| Figure S4A | *1151-Gal4* | *UAS-NΔECD, UAS-yellow-RNAi* | *GFP::CSL* |
| Figure S4A | *1151-Gal4* | *UAS-NΔECD, UAS-Mam-RNAi* | *GFP::CSL* |
| Figure S4B | *ms1096-Gal4, tub-Gal80ts* | *UAS-NICD[Δ1DR-PEST], UAS-LacZ* | *E(spl)-C[m∂ intA}, UAS-ParA::mCherry, UAS-NΔECD* |
| Figure S4B | *ms1096-Gal4, tub-Gal80ts* | *UAS-NICD[Δ1DR-PEST], UAS-Mam[DN]* | *E(spl)-C[m∂ intA}, UAS-ParA::mCherry, UAS-NΔECD* |
| Figure 5A-B | *ms1096-Gal4, tub-Gal80ts* | *UAS-NΔECD, GFP::Mam* | *mCherry::CSL,* |
| Figure 5C | *ms1096-Gal4, tub-Gal80ts* | *UAS-NΔECD,* | *GFP::CSL, E(spl)-C[m∂ intA}, UAS-ParA::mCherry* |
| Figure 5E, F, H, I | *1151-Gal4* | *UAS-CRYTEVc, UAS-OptIC-Notch{ω}::mCherry, Halo::Mam* | *GFP::CSL* |
| Figure S5A | *ms1096-Gal4, tub-Gal80ts* | *UAS-NΔECD, GFP::Mam* | *mCherry::CSL,* |
| Figure 6A | *1151-Gal4* | *UAS-LacZ, GFP::Mam* | *E(spl)-C[m∂ intA}, UAS-ParA::mCherry* |
| Figure 6A | *1151-Gal4* | *UAS-NΔECD, GFP::Mam* | *E(spl)-C[m∂ intA}, UAS-ParA::mCherry* |
| Figure 6B | *1151-Gal4* | *UAS LacZ, GFP::Rbp3* | *E(spl)-C[m∂ intA}, UAS-ParA::mCherry* |
| Figure 6B | *1151-Gal4* | *UAS-NΔECD, GFP::Rbp3* | *E(spl)-C[m∂ intA}, UAS-ParA::mCherry* |
| Figure 6C | *1151-Gal4* | *UAS-LacZ* | *E(spl)-C[m∂ intA}, UAS-ParA::mCherry, GFP::Med1* |
| Figure 6C | *1151-Gal4* | *UAS-NΔECD* | *E(spl)-C[m∂ intA}, UAS-ParA::mCherry, GFP::Med1* |

| | | | |
|---|---|---|---|
| Figure 6D | *1151-Gal4* | *UAS-NΔECD, hsp83-MCP::GFP* | *E(spl)-C[m∂ intA}, UAS-ParA::mCherry, E(spl)mβ-MS2* |
| Figure 6E | *1151-Gal4, Rbp1::mCherry* | *UAS-NΔECD* | *Halo::CSL, GFP::Med1* |
| Figure 6F | *1151-Gal4* | *UAS-NΔECD, GFP::Rbp3* | *E(spl)-C[m∂ intA}, UAS-ParA::mCherry* |
| Figure 6F | *1151-Gal4* | *UAS-NΔECD* | *E(spl)-C[m∂ intA}, UAS-ParA::mCherry, GFP::Med1* |
| Figure 6F | *1151-Gal4* | *UAS-LacZ, hsp83-MCP::GFP* | *E(spl)-C[m∂ intA}, UAS-ParA::mCherry, E(spl)mβ-MS2* |
| Figure 6F | *1151-Gal4* | *UAS-NΔECD, hsp83-MCP::GFP* | *E(spl)-C[m∂ intA}, UAS-ParA::mCherry, E(spl)mβ-MS2* |
| Figure S6A | *As Figure 6A* | | |
| Figure S6B | *As Figure 6B* | | |
| Figure S6C | *As Figure 6C* | | |
| Figure S6D | *As Figure 6D* | | |
| Figure S6E | *As Figure 6E* | | |
| Figure S6F-H | *As Figure 6F* | | |

**Table S4: p-values from statistical tests.**

| | | | |
|---|---|---|---|
| Figure 1B | Mam OFF vs. ON | n=32, 30 | p=1.4e-11 |
| Figure 1B | CSL OFF vs. ON | n=45, 28 | p = 1.4e-12 |
| Figure 1D | CSL OFF vs. ON | n=5, 7 | p=5.5e-3, |
| Figure 1D | Mam OFF vs. ON | n=5, 13 | p=2.3e-4 |
| Figure S1B | Mam ON wRi vs. ON HRi | n=25, 27 | p= 0.43 |
| Figure S1C | CSL ON wRi vs. ON HRi | n=38, 31 | p= 9.8e-06 |
| Figure 2A | ectopic array 12 vs. 48 | n=45, 40 | p=0.09 |
| Figure 2A | *E(spl)-C* 12 vs. 48 | n=45, 40 | p=0.75 |
| Figure 2C | NICD-PEST IDR Notch OFF vs. ON | n=62, 67 | p = 8.4e-03 |
| Figure 2C | CSL-nt IDR Notch OFF vs. ON | n=8, 40 | p=0.63 |
| Figure 2D | Nact FL vs. Nact ΔIDR,, | n=58, 55 | p=4.2e-4 |
| Figure S2C | CSL ct IDR Notch OFF vs. ON | n=39, 44 | p=0.69 |
| Figure S2C | NICD IDR Notch OFF vs. ON | n=83, 72 | p=3.4e-04 |
| Figure S2C | Med1 IDR Notch OFF vs. ON | n=32, 25 | p=0.04 |
| Figure S2C | Mam IDR Notch OFF vs. ON | n=18,36 | p=0.25 |
| Figure S2C | Mam nIDR Notch OFF vs. ON | n=12, 29 | p=0.14 |
| Figure 3A | DMSO vs. triptolide | n=49, 36 | p = 0.55 |
| Figure 3C | DMSO vs. A485 | n=47,62 | p=0.45 |
| Figure 3D | yRNAi vs CBP/p300 RNAi | n=26, 31 | p=0.17 |
| Figure 3E | Mam control vs Mam Med13Ri | n=30, 45 | p=9.8e-7 |
| Figure 3E | CSL control vs CSL Med13Ri | n=92, 36 | p=1.1e-4 |
| Figure 3F | Mam DMSO vs Mam Senexin B | n=27, 43 | p=1.2e-4 |
| Figure 3F | CSL DMSO vs CSL Senexin B | n=34, 37 | p=7.5e-5 |
| Figure 3G | ctrl vs. Cdk8 Ri, | n=33, 32 | p=9.4e-4 |
| Figure 3H | Notch OFF vs. ON | n=28, 31 | p=1.7e-10 |
| Figure S3A | DMSO vs. triptolide | n=18, 16 | p= 0.0118 |
| Figure S3A | DMSO vs. triptolide | n=9,8 | p= 0.17 |
| Figure S3B | DMSO vs. A485 | n=20, 6 | p= 2.9e-4 |
| Figure S3C | Cytoplasmic DMSO vs. A485 | n=32, 22 | p= 0.0081 |
| Figure S3C | *E(spl)-C* DMSO vs. A485 | n=13, 6 | p= 0.59 |
| Figure S3D | DMSO vs. Senexin A | n=27, 26 | p=7.5e-04 |
| Figure S3E | Control KD vs. Med13 KD | n=10, 11 | p= 0.004 |
| Figure S3F | DMSO vs. Senexin B | n=31, 28 | p= 0.71 |
| Figure S3G | DMSO vs. Senexin B | n=27, 31 | p= 4.8e-7 |
| Figure S3H | DMSO vs. NVP2 | n=30, 43 | p=0.83 |
| Figure S3I | DMSO vs. NVP2 | n=31, 14 | p=0.035 |
| Figure 4C | OFF vs. ON | n=28, 31 | p=1.7e-10 |

| | | | |
|---|---|---|---|
| Figure 4D’ | Cytoplasmic OFF vs. ON. | n=18, 20 | p=1.5e-3 |
| Figure 4D’ | Cytoplasmic ON ctrl vs. ON MamDN | n=20, 22 | p=8.1e-4 |
| Figure 4D’ | Cytoplasmic OFF vs. ON, MamDN | n=18, 22 | p=0.09 |
| Figure S4B | Control vs. MamDN | n=25, 28 | p=0.76 |
| Figure 5A | Mam 24 hrs ON vs 24 hrs ON, 4 hrs OFF | n=28, 19 | p=0.047 |
| Figure 5A | Mam 24 hrs ON, 8 hrs OFF vs 24 hrs ON, 4 hrs OFF | n=19, 21 | p = 8.9e-5 |
| Figure 5A | Mam 24 hrs ON, 8 hrs OFF vs 24 hrs ON, 4 hrs OFF | n=28, 21 | p = 2.5e-8 |
| Figure 5A | CSL 24 hrs ON vs 24 hrs ON, 4 hrs OFF | n=28, 19 | p = 0.10 |
| Figure 5A | CSL 24 hrs ON, 8 hrs OFF vs 24 hrs ON, 4 hrs OFF | n=19, 21 | p = 0.58 |
| Figure 5A | CSL 24 hrs ON, 8 hrs OFF vs 24 hrs ON, 4 hrs OFF | n=28, 21 | p = 0.22 |
| Figure 5C | CSL 24 hrs ON vs 24 hrs OFF | n=28, 21 | p = 0.42 |
| Figure 5C | Cytoplasmic 24 hrs ON vs 24 hrs OFF | n=30, 16 | p = 5.6e-3 |
| Figure 5C | Nuclear 24 hrs ON vs 24 hrs OFF | n=10, 5 | p = 0.14 |
| Figure 5F | Naïve vs. preactivated OFF | n=38, 21 | p = 8.6e-4 |
| Figure 5F | preactivated OFF vs. preactivated ON | n=21, 43 | p = 2.3e-5 |
| Figure 5F | Naïve vs. preactivated ON | n=38, 43 | p = 0.09 |
| Figure 5I | Naïve vs preactivated | n=11, 9 | p=1.7e-3 |
| Figure S5A | cytoplasmic 24 hrs ON vs 24 hrs ON, 4 hrs OFF | n=22, 30 | p = 0.04 |
| Figure S5A | cytoplasmic 24 hrs ON, 8 hrs OFF vs 24 hrs ON, 4 hrs OFF | n=30, 30 | p = 0.93 |
| Figure S5A | cytoplasmic 24 hrs ON, 8 hrs OFF vs 24 hrs ON, 4 hrs OFF | n=22, 30 | p = 0.06 |
| Figure S5A | nuclear 24 hrs ON vs 24 hrs ON, 4 hrs OFF | n=4, 4 | p = 0.49 |
| Figure S5A | nuclear 24 hrs ON, 8 hrs OFF vs 24 hrs ON, 4 hrs OFF | n=4, 4 | p = 0.73 |
| Figure S5A | nuclear 24 hrs ON, 8 hrs OFF vs 24 hrs ON, 4 hrs OFF | n=4, 4 | p = 0.73 |
